# A partial pathogenicity chromosome in *Fusarium oxysporum* is sufficient to cause disease and can be horizontally transferred

**DOI:** 10.1101/2020.01.20.912550

**Authors:** Jiming Li, Like Fokkens, Lee James Conneely, Martijn Rep

**Affiliations:** Molecular Plant Pathology, University of Amsterdam, Amsterdam, 1098 XH, the Netherlands

## Abstract

During host colonization, plant pathogenic fungi secrete proteins, called effectors, to facilitate infection. Collectively, effectors may defeat the plant immune system, but usually not all effectors are equally important for infecting a particular host plant. In *Fusarium oxysporum* f.sp. *lycopersici*, all known effector genes – also called *SIX* genes – are located on a single accessory chromosome which is required for pathogenicity and can also be horizontally transferred to another strain. To narrow down the minimal region required for virulence, we selected partial pathogenicity chromosome deletion strains by fluorescence-assisted cell sorting of a strain in which the two arms of the pathogenicity chromosome were labelled with *GFP* and *RFP*, respectively. By testing the virulence of these deletion mutants, we show that the complete long arm and part of the short arm of the pathogenicity chromosome are not required for virulence. In addition, we demonstrate that smaller versions of the pathogenicity chromosome can also be transferred to a non-pathogenic strain and they are sufficient to turn the non-pathogen into a pathogen. Surprisingly, originally non-pathogenic strains that had received a smaller version of the pathogenicity chromosome were much more aggressive than recipients with a complete pathogenicity chromosome. Whole genome sequencing analysis revealed that partial deletions of the pathogenicity chromosome occurred mainly close to repeats, and that spontaneous duplication of sequences in accessory regions is frequent both in chromosome deletion strains and in horizontal transfer (recipient) strains.

**Author Summary:** Fungal genomes can often be divided into a core genome, which is essential for growth, and an accessory genome which is dispensable. The accessory genome in fungi can be beneficial under some conditions. For example, in some plant-pathogenic fungi, virulence genes are present in the accessory genome, which enable these fungi to cause disease on certain hosts. In *Fusarium oxysporum* f.sp. *lycopersici*, which infects tomato, all host-specific virulence genes are located on a single accessory chromosome. This ‘pathogenicity chromosome’ can be horizontally transferred between strains. Here, we found that many suspected virulence genes are in fact not required for virulence because strains without a large part of the pathogenicity chromosome, including these genes, showed no reduced virulence. In addition, we demonstrate that partial pathogenicity chromosomes can be horizontally transferred to a non-pathogen. Surprisingly, originally non-pathogenic strains that had received a partial pathogenicity chromosome were more virulent than strains that had received the complete pathogenicity chromosome.

## Introduction

Accessory chromosomes, also called supernumerary chromosomes, B chromosomes, or lineage-specific chromosomes, were first discovered in *Hemiptera* in 1907 (1). However, it was only in 1991 that they were first reported in a fungus; the plant-pathogenic fungus *Nectria haematococca* (*Fusarium solani*) (2). Since then, accessory chromosomes have been found in more than 20 different species of fungi (3), including the plant pathogens *Fusarium oxysporum* (Fo) (4–7), *Fusarium solani* (8), and *Zymoseptoria tritici* (9–11). Accessory chromosomes are generally distinguished from core chromosomes by their relatively high number of repeats, lower gene density, distinct codon usage, different evolutionary trajectories and dispensability (12).

Although accessory chromosomes are dispensable, they can play an important role under specific conditions, such as conferring pathogenicity to specific plant species (3). For example, in *Alternaria*, host-selective toxin genes are located on accessory chromosomes which are responsible for causing disease on certain plant species (13–15). Recent findings in the hemibiotrophic plant pathogen *Colletotrichum higginsianum* showed that mutants without chromosome 11 are arrested during the biotrophic phase of infection (16). In contrast, loss of this chromosome had no clear effect on vegetative fitness, suggesting that this chromosome plays a specific role during infection (16). One of the most well-documented examples is the pathogenicity chromosome of Fo f.sp. *lycopersici* (Fol) (4,17,18). While this pathogenicity chromosome can be lost without affecting normal growth, strains without this pathogenicity chromosome cannot infect tomato plants (4,16). All 14 known effector genes (*<u>S</u>ecreted <u>I</u>n <u>X</u>ylem* genes, *SIX* genes) are located on this pathogenicity chromosome (18), and some of these effector genes were shown to contribute to virulence towards tomato plants, including *SIX1* (*AVR3*) (19), *SIX3* (*AVR2*) (20), and *SIX5* (21). Further studies on this pathogenicity chromosome showed that loss of (most of) the long arm (q arm) of this chromosome, including *SIX6*, *SIX9* and *SIX11*, did not significantly affect virulence (5).

Apart from conferring advantages in a certain environment, at least some accessory chromosomes can also be horizontally transferred from one strain to another (4,6,17,22). The first molecular evidence for horizontal chromosome transfer (HCT) in fungal plant pathogens was reported in *Colletotrichum gloeosporioides* (23). It was suggested that a 2-Mb chromosome in the biotype B isolate Bx most likely originated by a relatively recent transfer from biotype A. Shortly after, He and colleagues experimentally demonstrated horizontal transfer of a 2-Mb chromosome from biotype A to biotype B, however, no pathogenicity phenotype was transferred (22).

HCT has also been observed in Fo (4,6,17). When co-incubating a Fol strain with a non-pathogenic strain, the pathogenicity chromosome of Fol can be transferred to a non-pathogenic strain, turning the latter into a tomato-infecting strain (4,17). In some cases, a second accessory chromosome was co-transferred (4). Similarly, from Fo f.sp. *radicis-cucumerinum* (Forc), a single chromosome chr^RC^ can be transferred to a non-pathogen, turning the recipient into a cucurbit-infecting strain (6). The mechanisms behind HCT are largely unknown, but it is most likely that HCT happens through heterokaryosis, which was supported by the observation that transfer is not always restricted to accessory chromosomes, but a core chromosome (~4 Mb) could also be transferred (17).

In a previous study, we showed that the short arm (p arm) of the pathogenicity chromosome in Fol can be sufficient for causing disease on tomato plants (5). In order to test this hypothesis and narrow down the genes or regions that are essential for infection in Fol, we selected partial pathogenicity chromosome deletion strains. To achieve this, we inserted the *RFP* gene in the short arm (p arm) of the pathogenicity chromosome of a strain with the *GFP* gene on the q arm (16), and used fluorescence-assisted cell sorting to select spores without GFP or RFP (24). By testing the virulence of these deletion mutants, we show that less than half of the chromosome is sufficient for causing disease. In addition, we demonstrate that smaller versions of the pathogenicity chromosome can also be transferred to a non-pathogenic strain, with concomitant transfer of pathogenicity towards tomato.

## Results

### Construction of a Fol strain with *GFP* and *RFP* on either arm of the pathogenicity chromosome

To be able to select for partial pathogenicity chromosome deletion strains in Fol, we set out to create a strain with the *RFP* gene on the short arm (p arm) and the *GFP* gene on the long arm (q arm) of the pathogenicity chromosome. Fluorescence-assisted cell sorting (FACS) with this strain could then be used to select spores without either green fluorescence or red fluorescence. To construct this strain, the strain 14HG6B with *GFP* on the q arm of the pathogenicity chromosome was used as a starting point (5). To insert the *RFP* gene on the p arm, single copy genes *FOXG_14135* and *FOXG_16428* with relatively low expression during colonization of tomato plants (25) were selected for homologous recombination. With the additional purpose to investigate whether the *SIX10/12/7* gene cluster contributes to virulence, this gene cluster was also targeted for homologous recombination. The location of these genes is shown in Fig 1A.

**Fig 1:**
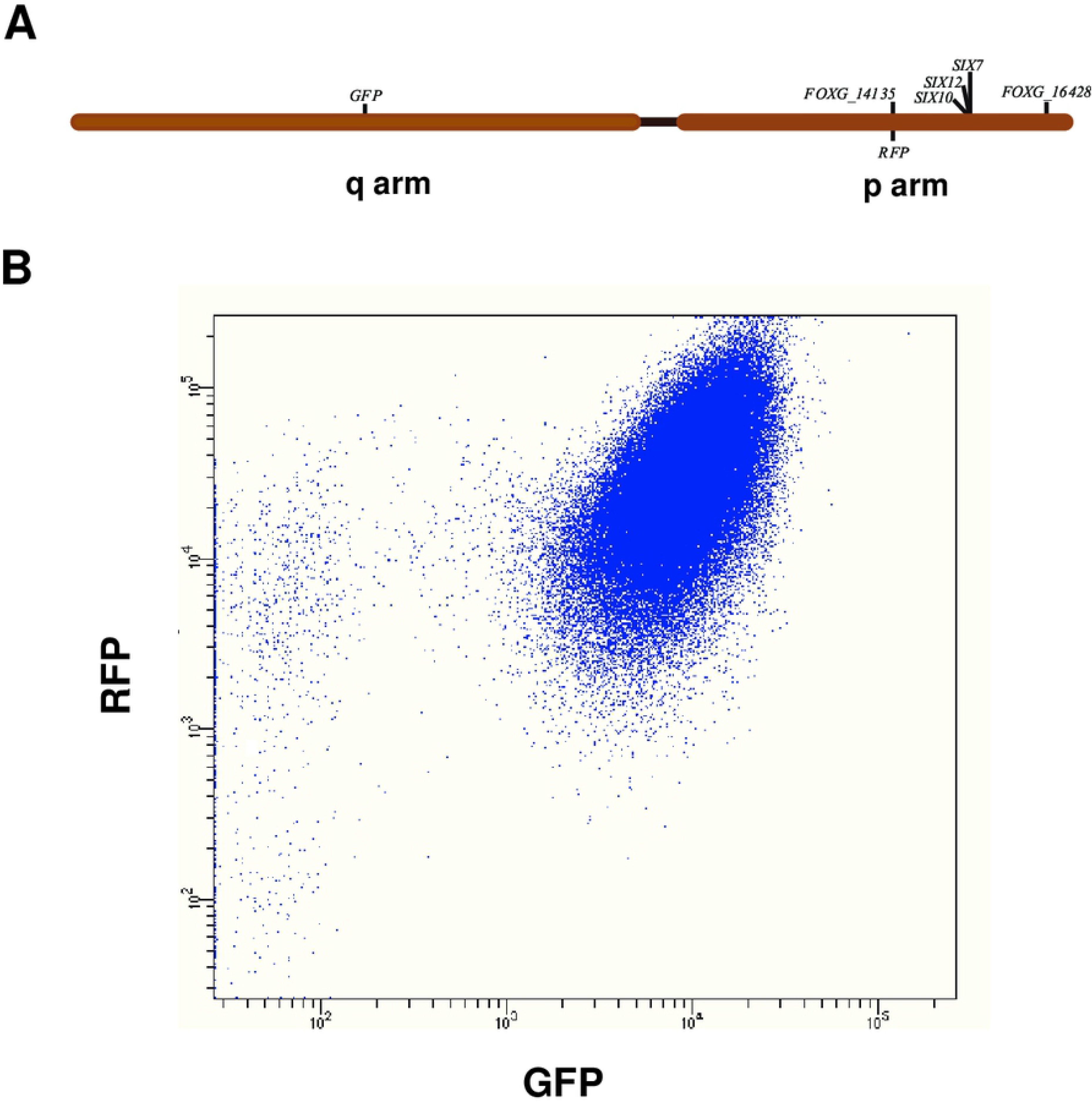
In cultures of Fol strain 14HGPR, loss of green fluorescence is much more frequent than loss of red fluorescence. (A) Schematic representation of the Fol pathogenicity chromosome and genes selected for replacement with *RFP*. The long arm is indicated as q arm, while the short arm is indicated as p arm. (B) Dot plot of a fluorescence assisted cell sorting experiment. Each blue dot represents a fungal spore. Most spores contain both red and green fluorescence. Some spores had lost green fluorescence, while very few spores had lost red fluorescence. Axis labels show the detection channel (X-axis: λ=488nm; Y-axis: λ=561nm).

To replace genes with *RFP*, *Agrobacterium*-mediated transformation was performed. For *SIX10/12/7*, no *in locus* transformant was found after checking 433 transformants in two rounds of transformation, but two spontaneous *SIX10/12* deletion mutants, 14HG6B_ΔSIX10_12#1 and 14HG6B_ΔSIX10_12#2, were found. Since these were derived from the same experiment, it could be that they are not independent but arose from a single event. In any case, *SIX10* and *SIX12*, as part of a ~20kb region, were lost in these strains, but *SIX7* was retained. In the effort to replace *FOXG_16428* with *RFP*, one *in locus* transformant, 14HG6B_ΔFOXG_16428, was obtained out of 945 transformants in three rounds of transformation. However, this transformant also contained (an) ectopic insertion(s) of the *RFP* construct. For the third gene, *FOXG_14135*, 300 transformants in two rounds of transformation were checked, and we found one *in locus* transformant without ectopic insertion, 14HG6B_ΔFOXG_14135 (called **14HGPR** from hereon). Fol transformations are summarized in Table S1. Strain 14HGPR was confirmed microscopically to have both red and green fluorescence, and *in locus* insertion was confirmed by PCR.

To assess the virulence of 14HGPR, bioassays were performed. Disease index and fresh weight were scored three weeks after inoculation, and no significant reduction of virulence was observed when comparing 14HGPR with the original strain 14HG6B and an ectopic control (Fig S1). Thus, 14HGPR was used for FACS experiments to obtain partial deletions of the Fol pathogenicity chromosome from both arms. In addition, bioassays were performed to assess virulence of 14HG6B_ΔSIX10/12#1, 14HG6B_ΔSIX10/12#2, and 14HG6B_ΔFOXG_16428, and no significant reduction in virulence was observed (data not shown).

### *GFP* fluorescence is much more frequently lost in spores of 14HGPR than *RFP* fluorescence

To obtain spontaneous deletions of the Fol pathogenicity chromosome from both arms, Fluorescence Assisted Cell Sorting (FACS) of strain 14HGPR was performed to select spores without green or red fluorescence. In total, 26 different cultures were started from single spore colonies in four different FACS experiments, and from these experiments 43 *GFP* deletion strains and 18 *RFP* deletion strains were kept for further analysis (Table S2).

The first FACS experiment served to determine the approximate rate of loss of green or red fluorescence in cultures of the 14HGPR strain. Six single colonies of 14HGPR were separately inoculated into NO_3_ medium (0.17% yeast nitrogen base, 3% sucrose, 100mM KNO_3_). After growing for five days, spore suspensions were obtained by filtering cultures through a double layer of mira-cloth and directly used for FACS. We observed that all the six cultures showed a similar pattern, with a large population of spores still containing both green and red fluorescence and a very small fraction without green or red fluorescence (Fig 1B). Strikingly, in all cultures more spores had lost green fluorescence than spores that had lost red fluorescence, as shown in Fig 1B. Only one single spore colony which had lost the *RFP* gene was kept for further analysis from this first experiment (Table S2).

For the second FACS experiment, 14HGPR was again mono-spored and five single colonies were transferred directly into NO_3_ medium and allowed to grow for five days. This time two hundred fifty spores without green fluorescence and 226 spores without red fluorescence were deflected on Potato Dextrose Agar (PDA) plates and allowed to form colonies (Table S3). Only 27 out of 250 (11%) colonies emerged on the plates on which spores without green fluorescence were deflected, while 199 spores selected for loss of red fluorescence (87%) formed colonies (Table S3). It turned out that 18 out of the 27 (66.7%) ‘red-selected’ colonies were confirmed to be GFP negative when checked by microscopy, but only three out of 199 (1.5%) ‘green-selected’ colonies were truly RFP negative (Table S3). Since spores without red fluorescence were extremely rare, the gating was set close to spores containing both red and green fluorescence, and this apparently resulted in many false negative spores. The details from all five cultures are shown in Fig S4. For the 18 ‘red’ and the three ‘green’ colonies, PCR was used to confirm loss of the *GFP* or *RFP* gene and other regions of the Fol pathogenicity chromosome (Table S2).

To obtain more independent deletion strains, a third FACS experiment was performed. With the aim of increasing the frequency of spontaneous loss of *RFP*, five single colonies of 14HGPR were grown on PDA plates for ten days before collecting and inoculating spores from plates into NO_3_ medium. After five days incubation at 25°C in the NO_3_ medium, the spores were transferred to 4°C for five days and 1 mL of these suspensions were transferred into new NO_3_ medium and allowed to grow for another seven days at 25°C before being subjected to sorting. Of spores without green fluorescence deflected, 74 out of 250 (30%) formed colonies, while of spores without red fluorescence deflected, 335 out of 375 (89%) formed colonies (Table S3). Fluorescence microscopy revealed that 63 out of the 74 (85%) ‘red’ colonies were truly green fluorescence negative. However, only ten out of 335 (3.0%) ‘green’ colonies were red fluorescence negative (Table S3). These 63 ‘red’ and ten ‘green’ colonies were also confirmed by PCR to have lost *GFP* or *RFP*, respectively. The details from all five cultures are shown in Fig S5. Twenty four out of 63 ‘red’ and nine out of ten ‘green’ deletion strains were further checked for loss of other regions of the pathogenicity chromosome (Table S2). Concluding, using a longer culturing regime including incubation at 4°C, an increase in the frequency of loss of *GFP* was observed for all the five cultures, but no significant increase in the frequency of loss of *RFP* was observed when compared with the second FACS experiment (Table S3, Table S4 and Table S5).

So far, we obtained a large variety of partial deletions from the q arm of the pathogenicity chromosome, but from the p arm only 13 partial deletion strains with limited variation were found (Table S2). In a final attempt to obtain more partial deletions from the p arm of the pathogenicity chromosome, a fourth FACS experiment was performed to only select spores without red fluorescence. In this case, ten single colonies of 14HGPR were grown on PDA plates for one month at 25°C, then the spores were collected from the plates and inoculated into NO_3_ medium. The cultures were incubated for five days in NO_3_ medium before being used for FACS. Out of 246 single colonies growing from deflected ‘green’ spores, only five (2%) had truly lost red fluorescence when checked microscopically (Table S3). PCR of these five single colonies confirmed that the *RFP* gene was lost in all cases (Table S2). The details from all ten cultures are shown in Fig S6.

### Illumina whole genome sequencing confirms partial deletions of the Fol pathogenicity chromosome and reveals multiplications

To more accurately assess which sequences of the pathogenicity chromosome had been lost and whether changes had also occurred in other parts of the genome, we selected ten deletion strains with different deletion patterns for Illumina whole genome sequencing (Table S7). Of these ten deletion strains, six strains had lost part of the q arm, and four strains had lost part of the p arm. To determine sequence changes in the deletion strains, Illumina short-read mapping of both *GFP* deletion strains and *RFP* deletion strains was performed to the SMRT assembly of Fol4287 (Fig 2A and Fig 3A; Fig S2 and Fig S3). In addition, genome sequence reads from three previously obtained deletion strains, named 14-2, 14-4 and 14-7 (5) were mapped to the newly generated SMRT assembly of Fol4287 (Fig 2A and Fig S2). As a reference, Fol4287 Illumina sequencing reads were retrieved from SRA and were also mapped (Fig 2A and Fig 3A; Fig S2A and Fig S3A).

**Fig 2:**
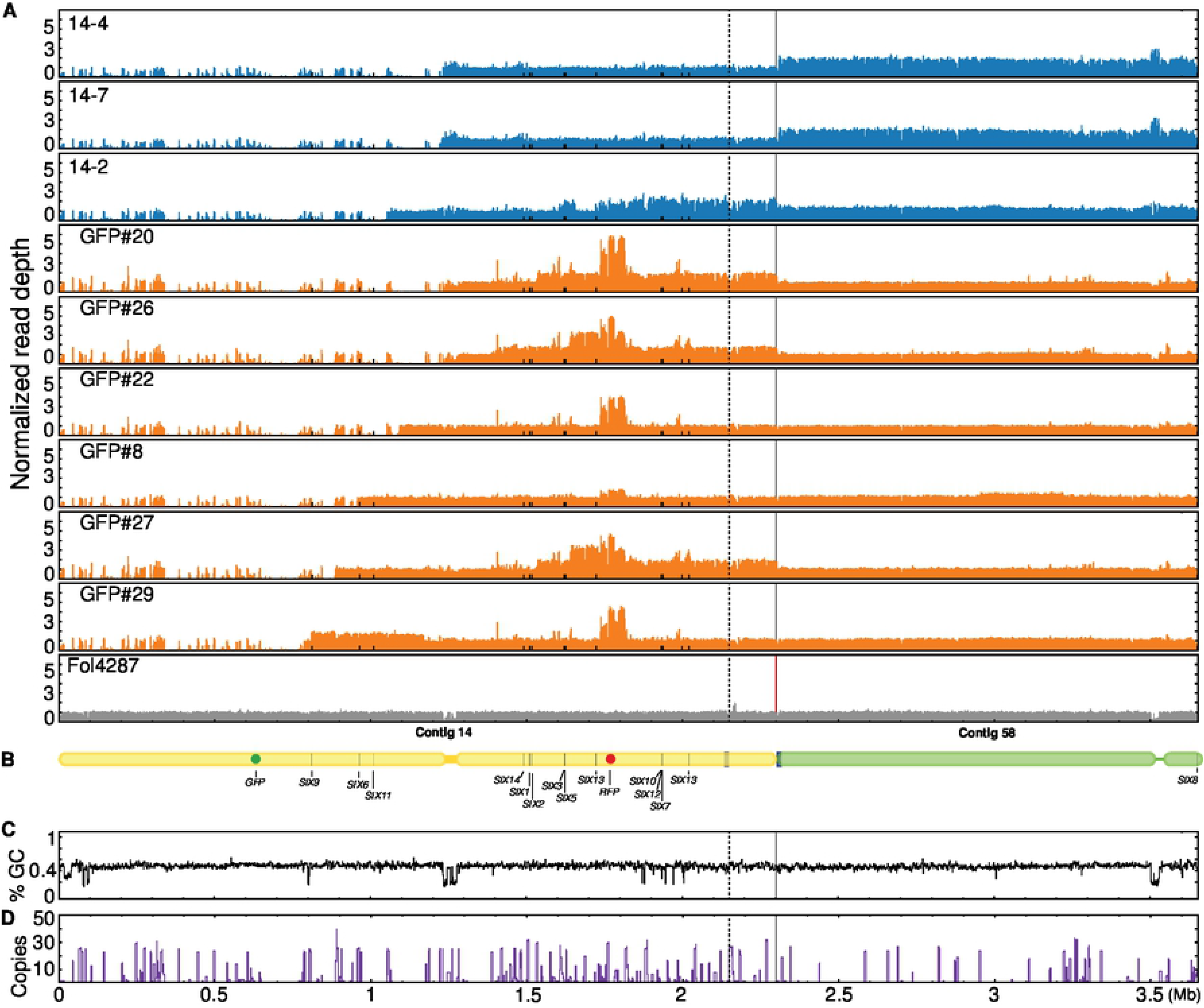
Illumina read mapping confirms partial deletions and reveals multiplications in the Fol pathogenicity chromosome in *GFP* deletion strains. (A) Reads of nine *GFP* deletion strains were mapped to the SMRT assembly of Fol4287. As reference, Illumina reads of Fol4287 itself was also mapped. For comparison of differences within and between deletion strains, all genome coverage was normalized. All deletion strains had lost part of or the complete q arm of the pathogenicity chromosome. In addition, multiplications had occurred in the remaining part of the pathogenicity chromosome or contig 58 in some deletion strains. Part of contig 58 belongs to the pathogenicity chromosome as indicated between the solid and dotted lines. (B) Schematic representation of the pathogenicity chromosome (contig 14 and part of contig 58) and, for comparison, the rest of contig 58. Secreted In Xylem (*SIX*) genes are also indicated. GC content (C) and repeat distribution across the genome (D) are also displayed.

**Fig 3:**
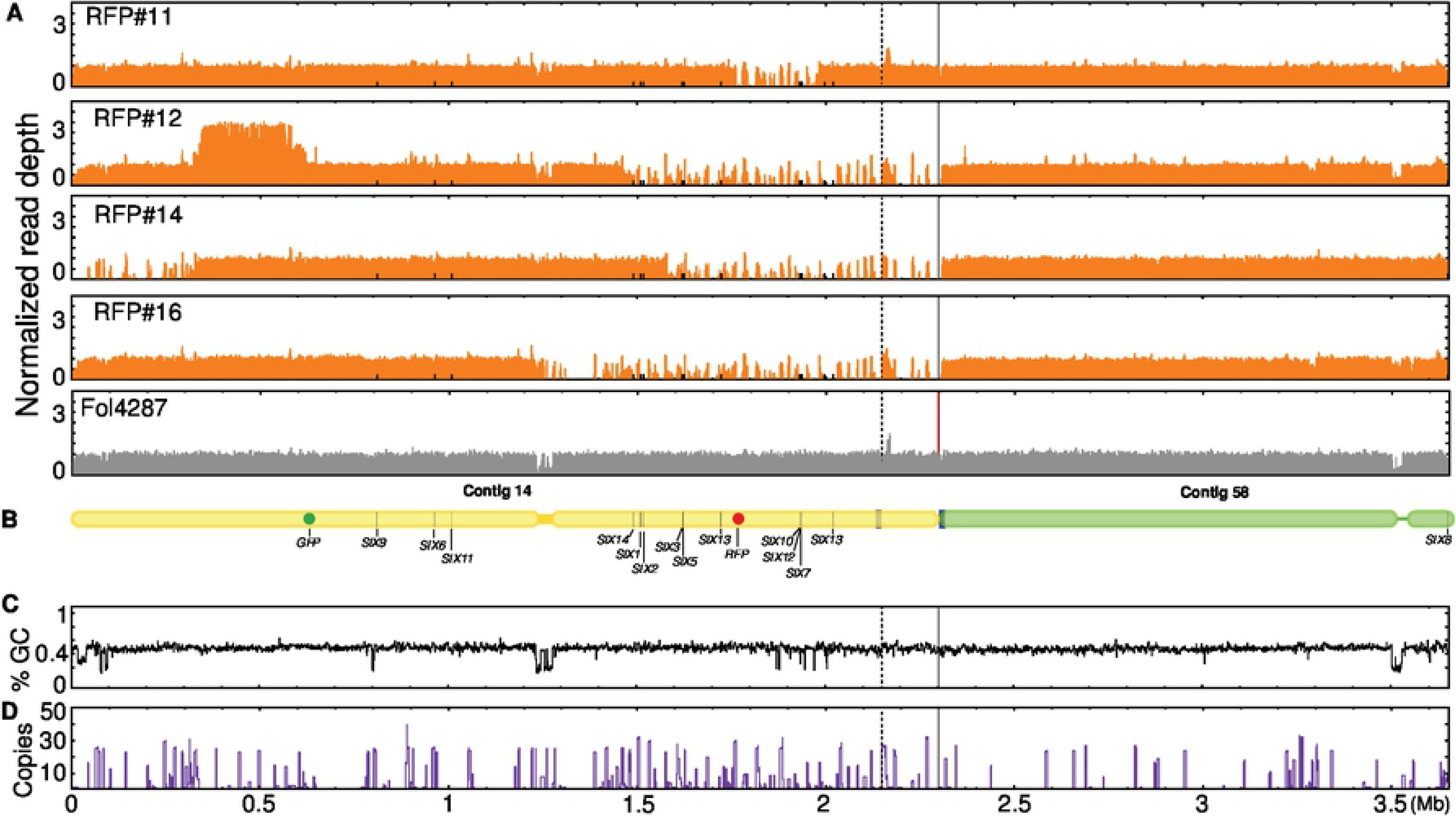
Illumina read mapping confirms partial deletions and reveals multiplications in the Fol pathogenicity chromosome in *RFP* deletion strains. (A) Reads of four *RFP* deletion strains were mapped to the SMRT assembly of Fol4287. As reference, Illumina reads of Fol4287 itself was also mapped. For comparison of differences within and between deletion strains, all genome coverage was normalized. All deletion strains had lost part or complete p arm of the pathogenicity chromosome. Multiplications had occurred only in ΔRFP#12. Surprisingly, the end of the q arm was lost in ΔRFP#14. Part of the contig 58 belongs to the pathogenicity chromosome as indicated between the solid and dotted lines. (B) Schematic representation of the pathogenicity chromosome (contig 14 and part of contig 58) and, for comparison, the rest of contig 58. Secreted In Xylem (*SIX*) genes are also indicated. GC content (C) and repeat distribution across the genome (D) are also displayed.

All nine *GFP* deletion strains were confirmed to have lost part or complete q arm of the pathogenicity chromosome (Fig 2A). Strains that had completely lost the q arm of the pathogenicity chromosome include 14-4, 14-7, ΔGFP#20 and ΔGFP#26. These deletions had happened close to or in the centromere region of the pathogenicity chromosome. Three *SIX* genes are located on the q arm, *SIX9*, *SIX6* and *SIX11*, which were lost in all these four deletion strains. 14-4 and 14-7 were probably derived from the same deletion event since these two deletion strains showed exactly the same read density pattern (Fig. 2A). The deletion strains 14-2 and ΔGFP#22 still contain a small part of the q arm, but no *SIX* genes are present in this part. The truncation of the pathogenicity chromosome in strain ΔGFP#8 is quite close to *SIX6*, and *SIX6* was lost in this strain (Table S2). *SIX6* and *SIX11* are present in the deletion strain ΔGFP#27, while *SIX9*, *SIX6* and *SIX11* are all present in ΔGFP#29.

In addition to partial or complete deletion of the q arm, multiplication of certain regions of the remaining part of the pathogenicity chromosome had also occurred for seven out of nine *GFP* deletion strains (Fig 2A). For all newly obtained deletion strains in this study, multiplication had occurred in the region where *RFP* was inserted, and this probably happened during insertion of *RFP* into this location. Part of the population of ΔGFP#8 used to prepare genomic DNA for sequencing probably had lost the multiplication in the *RFP* region as lower read densities were observed in this region. Duplication of the q arm was only observed for ΔGFP#29. Except multiplication of the *RFP* region, other large multiplications of the p arm of the pathogenicity chromosome were observed for deletion strains 14-2, ΔGFP#20, ΔGFP#26 and ΔGFP#27. Surprisingly, the whole contig 58 was duplicated in the deletion strains 14-4 and 14-7.

Lastly, to further assess whether deletions or multiplications could be linked to repeats, the distribution of repeats of the pathogenicity chromosome were determined (Fig 2D). Except for the deletion in strain ΔGFP#22, the remaining eight deletions had occurred close to repeats (Fig 2A and D). Among them, four deletions had occurred in the centromeric region, and the other four deletions had occurred close to repeats in different locations. No large changes were observed in the core genome of the *GFP* deletion strains (Fig S2). Interestingly, however, four out of the six newly generated deletion strains of the q arm showed the same deletion in contig 47. Additional deletions and duplications in the core genome were only observed for ΔGFP#29, including a relatively large deletion at the end of contig 3, a smaller deletion at the end of contig 7, and a duplication at the end of contig 61.

The four *RFP* deletion strains all showed different deletion patterns (Fig 3A). Strain ΔRFP#11 had only lost a small region of the p arm of the pathogenicity chromosome, including the *SIX10/SIX12/SIX7* gene cluster. The end of the p arm of this deletion strain was still present. Strain ΔRFP#12 had lost a larger part of the p arm and this lost region including *SIX14*, *SIX1*, *SIX2*, *SIX3, SIX5*, *SIX13*, and *SIX10/12/7*. Interestingly, ΔRFP#14 not only had lost part of the p arm, but it also had lost the end of the q arm. In this deletion strain, only *SIX14*, *SIX1* and *SIX2* are still present on the p arm of the pathogenicity chromosome. Complete loss of the p arm of the pathogenicity chromosome was observed for ΔRFP#16. In contrast to the common multiplications observed for most *GFP* deletion strains, multiplications were only observed for one *RFP* deletion strain, ΔRFP#12.

In the *RFP* deletion strains, all deletions and multiplications had occurred close to repeats (Fig 3A and D). Surprisingly, all four *RFP* deletion strains had the same deletion in contig 47 as the *GFP* deletion strains (Fig S3A). Moreover, strain ΔRFP#12 contains one relatively large deletion at the end of contig 2 (Fig S3A and C).

To conclude, partial deletions of the pathogenicity chromosome were confirmed for all the deletion strains. In addition, multiplications and additional deletions were also observed on the pathogenicity chromosome as well as other parts of the genome in some strains. Lastly, almost all deletions and multiplications had occurred in or close to repetitive regions.

### A partial Fol pathogenicity chromosome can be transferred to a non-pathogenic strain

To test which parts of the Fol pathogenicity chromosome can be horizontally transferred, chromosome transfer experiments were performed (26) by co-incubating each of the selected 24 deletion strains containing different partial deletions with hygromycin or zeocin-resistant transformants of non-pathogenic strain Fo47 (‘recipient strains’) in five independent experiments (Table S8). Since the recipient strains produced more spores than the donor strains, we decided to co-incubate the donor strains and the recipient strains in different ratios, including 1:1, 10:1, and 20:1. Chromosome transfer was observed when using ratios of 1:1 or 10:1. Since the transfer frequency was extremely low, no significant difference in transfer frequency between these ratios could be determined. Co-incubation of donor and recipient strains was performed on Potato Dextrose Agar (PDA) medium or Czapek Dox Agar (CDA) in two different experiments (Table S9). Again, no significant difference in transfer frequency was observed. Since Shahi *et al*. (2016) showed that CAT medium (0.17% YNB, 25mM KNO_3_) facilitates heterokaryon formation, which could result in horizontal chromosome transfer, we also co-incubated the donor and recipient strains in CAT medium for three days before plating spores on PDA or CDA plates in one of the HCT experiments. However, no successful transfer events were observed (Table S9, HCT_IV).

Through these five experiments, we identified four strains, ΔGFP#8, ΔGFP#26, ΔGFP#29, and ΔRFP#1, with the ability to transfer its partial pathogenicity chromosome to Fo47 (Table S8). Horizontal transfer of partial chromosomes was confirmed by PCR using primers specific to the recipient strains and primers targeting different parts of the pathogenicity chromosome (Table S10). For donor strains ΔGFP#26 and ΔRFP#1, seven double drug-resistant colonies were found for each, designated HCT_ΔGFP#26-1 to -7 and HCT_ΔRFP#1-1 to -7. For donor strains ΔGFP#8 and ΔGFP#29, ten and six double drug-resistant colonies were obtained, designated HCT_ΔGFP#8-1 to -10 and HCT_ΔGFP#29-1 to -6. Among these donor strains, ΔGFP#8, ΔGFP#26, and ΔGFP#29 had lost different parts of the q arm, while ΔRFP#1 had lost a small part of the p arm. Transfer of partial chromosomes with large deletions of the p arm were not obtained, despite several attempts (Table S8).

It was observed earlier that chromosome size can change during horizontal chromosome transfer (6,17). To assess karyotypes of HCT-strains and donor strains, CHEF gel analysis was performed (Fig 4). One progeny strain from each donor strain (HCT_ΔRFP#1-7, HCT_ΔGFP#29-2, HCT_ΔGFP#8-2, and HCT_ΔGFP#26-1) was selected. As expected, all HCT-strains showed the karyotype of the recipient strain (Fo47) with an extra chromosome. For HCT_ΔRFP#1-7 and HCT_ΔGFP#8-2, the size of the extra chromosome is similar to the presumed partial pathogenicity chromosome in donor strains ΔRFP#1 and ΔGFP#8, respectively, which is consistent with the PCR results (Table S10) and the sequencing data (Fig 2A). However, in donor strain ΔGFP#29, instead of an expected smaller version of the pathogenicity chromosome as suggested by the PCR and sequencing data (Table S10 and Fig 2A), an extra chromosome of around 4 Mb was observed, suggesting translocation of the remaining part of the pathogenicity chromosome to another chromosome. After horizontal chromosome transfer from ΔGFP#29 to Fo47, an even larger chromosome (around 5 Mb) was found in the background of Fo47. In donor strain ΔGFP#26, which had lost the whole q arm of the pathogenicity chromosome (Fig 2A), a larger version (~2.5 Mb) of the pathogenicity chromosome was observed, which can be explained by the multiplication of the remaining part of the pathogenicity chromosome (Fig 2A). This ~2.5Mb chromosome apparently became smaller after horizontal chromosome transfer (around 2 Mb in HCT_ΔGFP#26-1).

**Fig 4:**
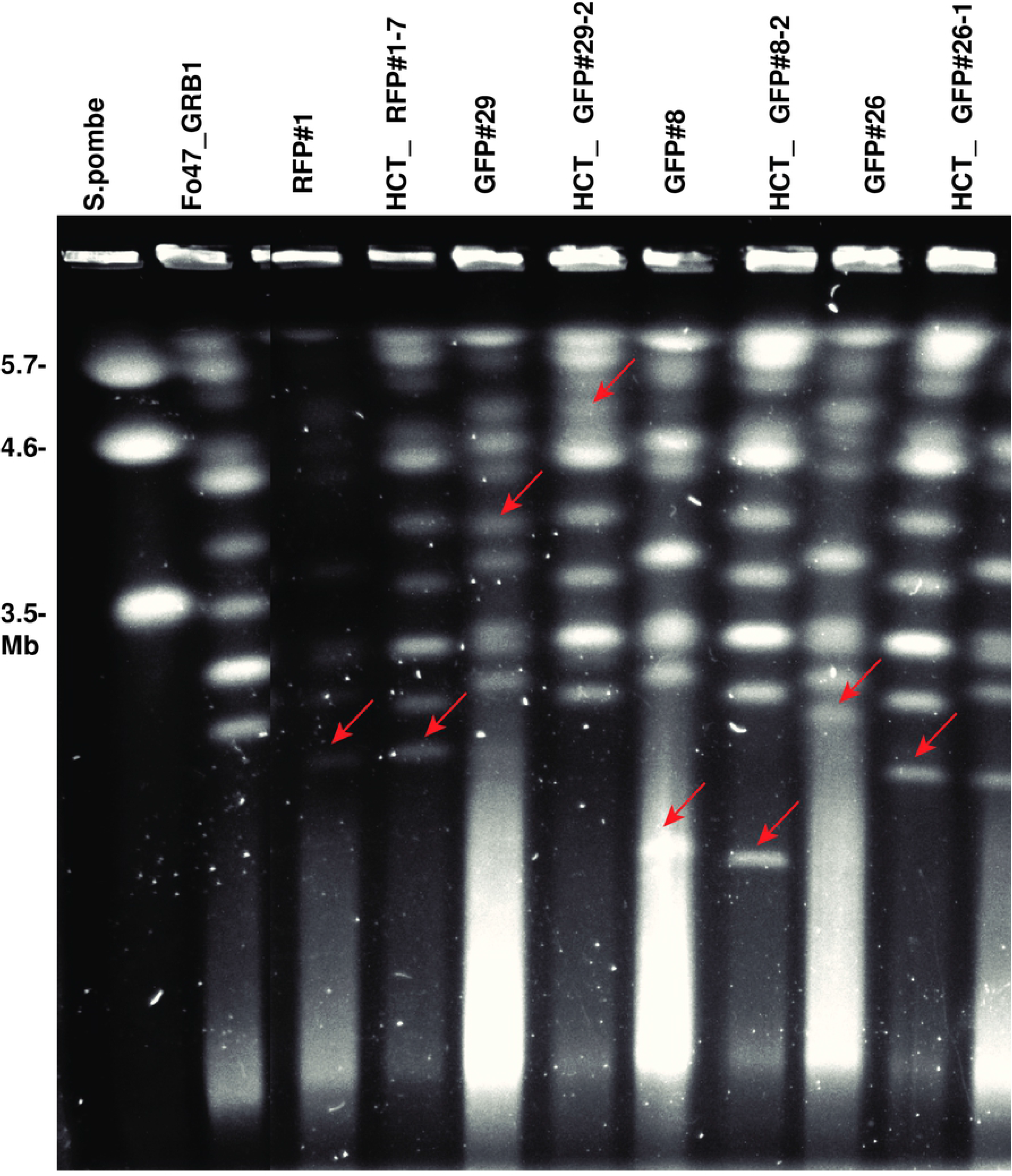
Contour-clamped homogeneous electric field (CHEF) electrophoresis confirms horizontal chromosome transfer. HCT-strains HCT_ΔRFP#1-7, HCT_ΔGFP#29-2, HCT_ΔGFP#8-2, and HCT_ΔGFP#26-1 all showed the karyotype of the recipient strain (Fo47), with an extra chromosome indicated with a red arrow. The extra chromosome in HCT_ΔRFP#1-7 and HCT_ΔGFP#8-2 is of a size similar to that of the partial pathogenicity chromosome in the donor strains ΔRFP#1 and ΔGFP#8 (red arrows), respectively. However, the extra chromosome in HCT_ΔGFP#29-2 and HCT_ΔGFP#26-1 is of a different size compared to the extra chromosome in donor strain ΔGFP#29 and ΔGFP#26, respectively (red arrows). Chromosomes of *S. pombe* was used as a marker. The figure was cropped.

### Illumina whole genome sequencing confirms transfer of partial pathogenicity chromosomes and reveals multiplications during chromosome transfer

To identify the sequences involved in karyotype changes (observed from the CHEF gel) during horizontal chromosome transfer, whole genomes of the HCT-strains HCT_ΔGFP#29-2, HCT_ΔGFP#8-2 and HCT_ΔGFP#26-1 were sequenced. To identify which sequences were newly acquired during horizontal chromosome transfer, stringent Illumina short-read mapping of HCT-strains against the SMRT assembly of Fol4287 was performed (Fig 5 and Fig S4). As reference, Illumina reads of donor strains were also mapped. As shown in Fig S4, and in accordance with the karyotype patterns as observed in the CHEF gel, all HCT-strains had the background of Fo47 with some extra sequences from the respective donor strains. Remarkably, in HCT-strain HCT_ΔGFP#29-2, the partial pathogenicity chromosome was almost fully duplicated after horizontal chromosome transfer, and contig 58 was co-transferred (Fig 5 and Fig S4). This contig 58 corresponds to the co-transferred chromosome in Fol007 (4).

**Fig 5:**
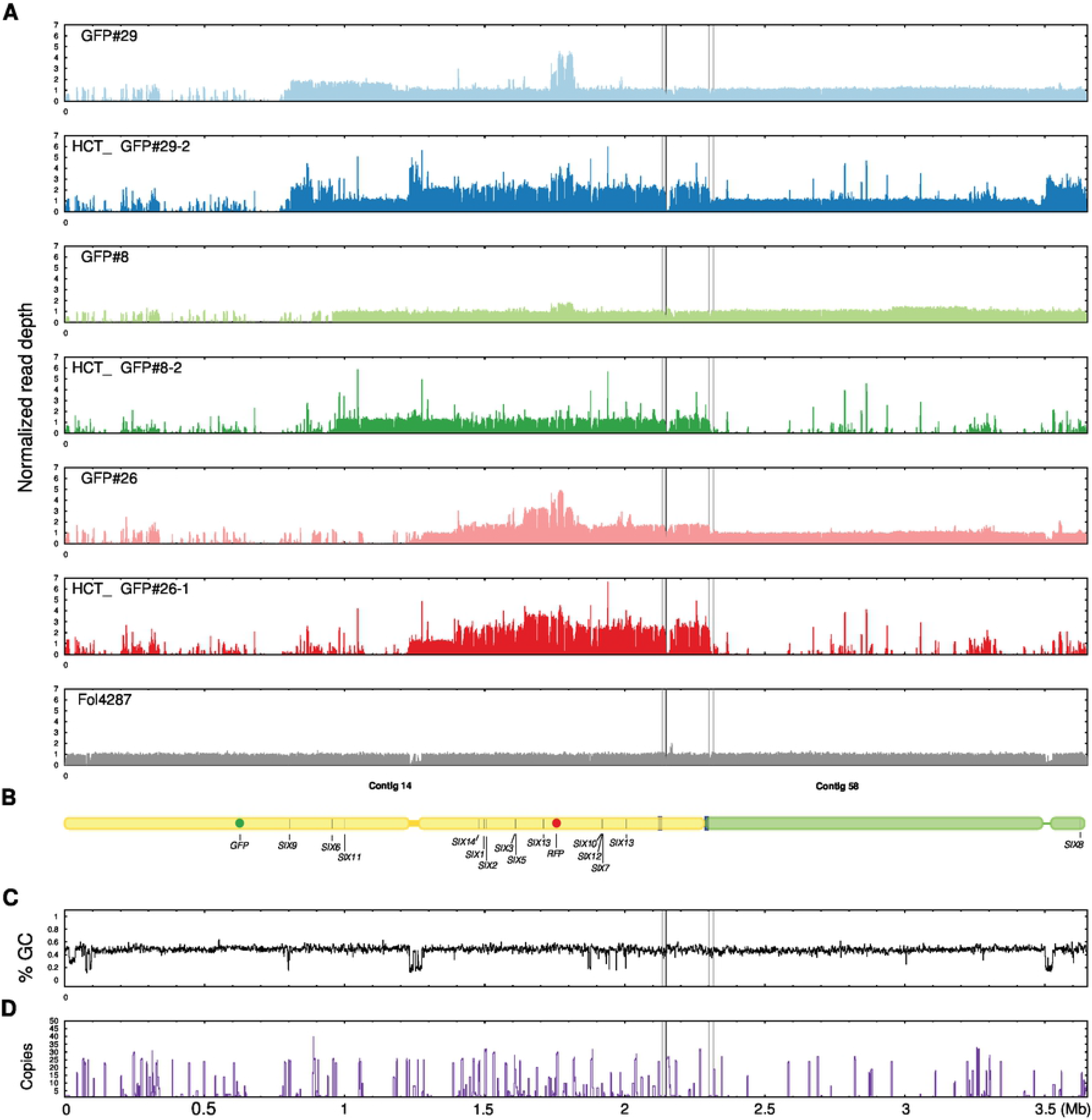
Stringent mapping of Illumina reads of HCT strains and donor strains to the SMRT assembly of Fol4287 confirms partial pathogenicity chromosome transfer. (A) Illumina reads of three HCT strains (HCT_ΔGFP#29-2, HCT_ΔGFP#8-2 and HCT_ΔGFP#26-1) and their respective donor strains (ΔGFP#29, ΔGFP#8 and ΔGFP#26-1) were mapped to the SMRT assembly of Fol4287, and only those reads that mapped completely and without any mismatches were selected. As reference, Illumina reads of Fol4287 were also mapped. Partial pathogenicity chromosome transfer was confirmed for all HCT strains. Surprisingly, in HCT_ΔGFP#29-2, co-transfer contig 58 was observed while this was not observed for the other two HCT strains. In HCT_ΔGFP#29-2, large multiplications of the remaining part of the pathogenicity chromosome and the end of contig 58 (not part of the pathogenicity chromosome) were also observed. Sequence multiplication during horizontal chromosome transfer was also observed in HCT_ΔGFP#26-1, but not in HCT_ΔGFP#8-2. (B) Schematic representation of the pathogenicity chromosome (contig 14 and part of contig 58) and, for comparison, the rest of contig 58. Location of *SIX* genes and *GFP* and *RFP* are indicated. GC content (C) and repeat distribution across the genome (D) are also displayed.

Surprisingly, in the same strain a large part of the accessory contig 47 was also transferred (Fig S4). This could explain the large chromosome band observed in the CHEF gel (Fig 4). Consistent with the CHEF gel, HCT_ΔGFP#8-2 had received the partial pathogenicity chromosome from the donor strain ΔGFP#8, and no deletions or multiplications were observed after horizontal chromosome transfer. The partial pathogenicity chromosome from the donor strain ΔGFP#26 were fully transferred to the HCT-strain HCT_ΔGFP#26-1, but differences in multiplication were observed between ΔGFP#26 and HCT_ΔGFP#26-1. These differences could explain the chromosome size difference on the CHEF gel (Fig 4). No core chromosome transfer was observed in any HCT strain (Fig S4).

### Single chromosome sequencing confirms partial pathogenicity chromosomes in both donor and HCT strains

To further confirm that we correctly identified the putative pathogenicity chromosomes in the CHEF gel, we cut the putative pathogenicity chromosome bands from the gel and isolated DNA from the gel pieces for sequencing. In total, eight strains were selected, including three donor *GFP* deletion strains (ΔGFP#29, ΔGFP#8, ΔGFP#26), three HCT strains (HCT_ΔGFP#29, HCT_ΔGFP#8, HCT_ΔGFP#26), and two *RFP* deletion strains (ΔRFP#11 and ΔRFP#12). First, a CHEF gel was run (Fig S5), bands were cut from the gel and 11 samples (Table S7) were sent for Illumina sequencing. Reads obtained from each sample were mapped to the SMRT assembly of Fol4287 (Fig S6). Most bands were successfully sequenced and contained sequences from the pathogenicity chromosome, as expected (Fig S6). For example, band ΔGFP#8_SC in donor strain ΔGFP#8 and band HCT_ΔGFP#8_SC in the recipient strain HCT_ΔGFP#8 were confirmed to both contain the partial pathogenicity chromosome. Similarly, the bands in the donor strain ΔGFP#26 and the recipient strain HCT_ΔGFP#26 were also confirmed to contain the partial pathogenicity chromosome. For the third pair of donor and recipient strain, ΔGFP#29 and HCT_ΔGFP#29-2, we observed three extra bands in the donor (ΔGFP#29) and two extra bands in the recipient strain (HCT_ΔGFP#29-2). Therefore, five bands were cut and the isolated DNA sequenced. Probably because of low DNA yield from the smallest band, ΔGFP#29_SC_XS, no reads of this sample could be mapped to the SMRT assembly. For the other two bands from the donor strain, ΔGFP#29_SC_L contained the partial pathogenicity chromosome, while ΔGFP#29_SC_S contained sequences from contig 7 and part of the pathogenicity chromosome. It is most likely, therefore, that the partial pathogenicity chromosome was partially duplicated and translocated to core contig 7. For the corresponding recipient strain HCT_ΔGFP#29-2, the extra band HCT_ΔGFP#29_SC_L contained sequences of the pathogenicity chromosome as well as contig 58, and part of contig 47, and all these sequences were originated from the donor strain. This is consistent with the whole genome mapping data, which showed that these sequences were transferred (Fig S4). For the second band in the recipient strain, HCT_ΔGFP#29_SC_S, reads mapped abundantly to core contig 5 and much fewer reads mapped to the pathogenicity chromosome, which we suspect to be background. Finally, the bands ΔRFP#11_SC and ΔRFP#12_SC from *RFP* deletion strains ΔRFP#11 and ΔRFP#12, respectively, were confirmed to contain the expected partial pathogenicity chromosome (Fig S6).

### A partial pathogenicity chromosome is sufficient to cause disease on tomato

To investigate which parts of the pathogenicity chromosome of Fol are required for virulence, twenty-two deletion strains (Fig 6) with different deletions in either arm of the pathogenicity chromosome were selected to assess pathogenicity. In addition, strains 14-2 and 14-7 obtained earlier, with large deletions in the q arm of the pathogenicity chromosome, and showing no reduced virulence on tomato in an earlier investigation, were included as controls (5). Again, we did not observe reduced virulence with these strains. Consistently, deletion strain ΔGFP#26, generated in this study and with complete loss of the q arm of the pathogenicity chromosome (Fig 2), showed no reduced virulence compared to the parental strain, 14HGPR. We conclude from this that the entire q arm is not required for virulence under the tested conditions. Strain ΔRFP#11, which had lost part of the p arm of the pathogenicity chromosome (Fig 3), including the *SIX10/12/7* gene cluster, also showed no reduced virulence, suggesting that this part of the chromosome is also not required for virulence. In contrast, strain ΔRFP#14, with a larger deletion of the p arm, did not show any virulence. Compared to ΔRFP#11, *SIX3*, *SIX5* and *SIX13* were lost in ΔRFP#14. Strains ΔRFP#12 and ΔRFP#16, which had lost an even larger part of the p arm, also could not cause any disease on tomato plants. Since *SIX3* and *SIX5* have been shown to contribute to contribute to virulence (20,21), we transformed these to genes together to strain ΔRFP#14, but this did not lead to regaining of virulence (results not shown).

**Fig 6:**
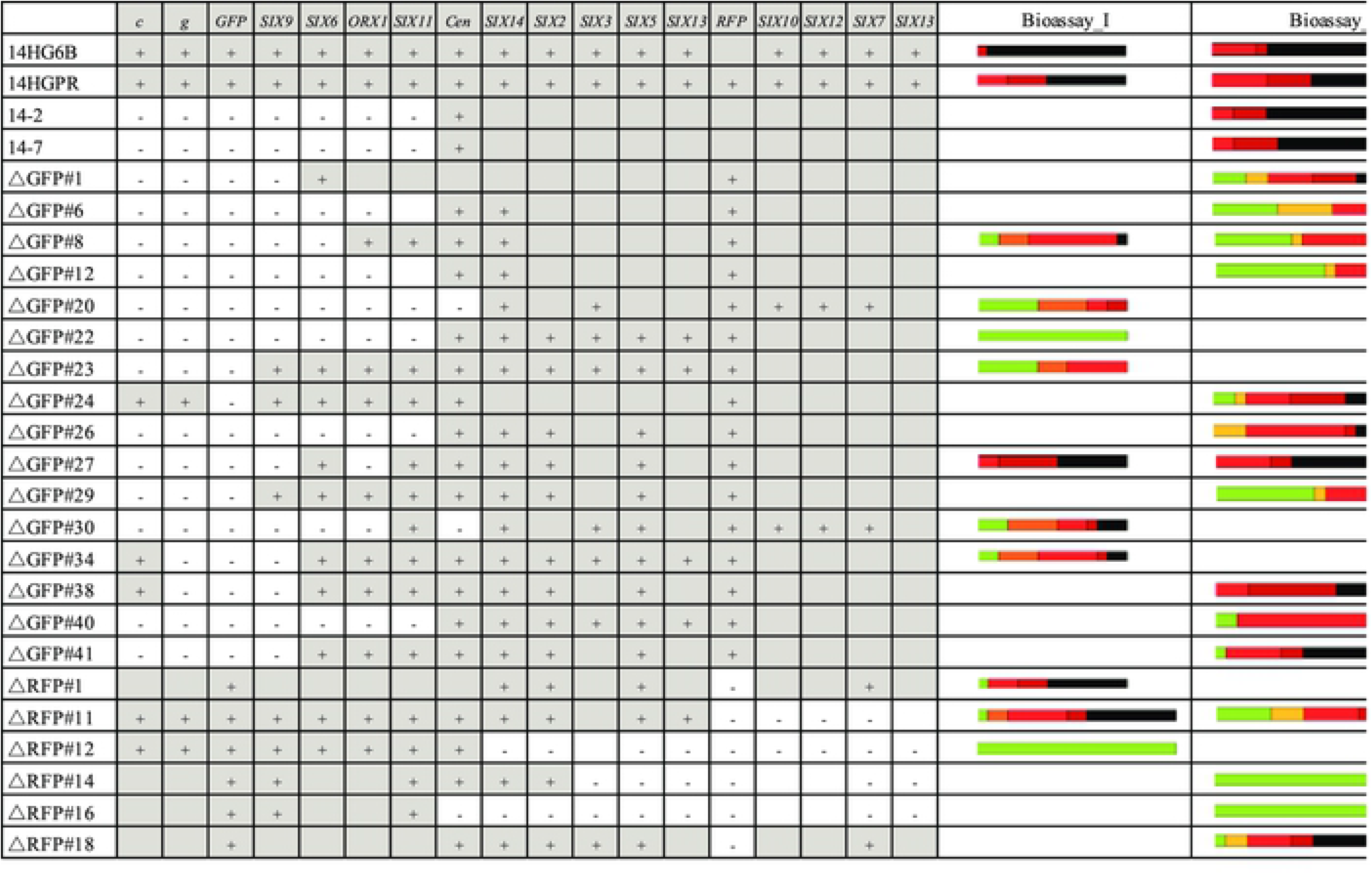
A partial pathogenicity chromosome in Fol is sufficient to cause disease on tomato plants. Bioassays were performed to assess virulence of deletion strains. Ten days old tomato seedlings were inoculated with 1*10^7^ spores/mL at 25°C, and disease index (DI) of infected tomato plants was scored three weeks after inoculation. 14HGPR showed no reduced virulence compared to the parental strain 14HG6B. Deletion strains 14-2, 14-7 and ΔGFP#26, with almost complete loss of the q arm, showed no reduced virulence. Similarly, ΔRFP#11 without the *SIX10/12/7* gene cluster showed no reduced virulence. Deletion strains with a larger deletion of the p arm (ΔRFP#12, ΔRFP#14 and ΔRFP#16) did not cause disease on tomato plants. Disease index was scored on a scale of 0–4 (0, no symptoms; 1, one brown vessel below the cotyledons; 2, one or two brown vascular bundles at cotyledons; 3, three brown vascular bundles and growth distortion; 4, all vascular bundles are brown, plant either dead or very small and wilted). Kruskal-Wallis test was performed on disease index.

### A partial pathogenicity chromosome can turn an endophyte into a pathogen

To investigate whether the endophytic strain Fo47 becomes a pathogen on tomato plants after receiving a partial Fol pathogenicity chromosome, three HCT-strains derived from each donor strain were selected to assess their virulence. For the four donor strains, ΔRFP#1, ΔGFP#29, ΔGFP#8 and ΔGFP#26, there was no significant difference in virulence compared to the parental strain 14HGPR (Fig 7). Fo47 could not cause any disease symptoms on tomato plants. However, all the HCT-strains were pathogenic to tomato plants with some variations in Disease Index (DI). For HCT-strains derived from ΔRFP#1, HCT_ΔRFP#1-5, HCT_ΔRFP#1-6, and HCT_ΔRFP#1-7, which acquired an almost complete pathogenicity chromosome, a relatively low disease index was observed (Fig 7 and Fig 8), which is consistent with the results from previous studies (4,5). Surprisingly, for HCT_ΔGFP#29-1, HCT_ΔGFP#29-2 and HCT_ΔGFP#29-3, much higher virulence was observed, comparable to, or even stronger than, the donor strain ΔGFP#29 (Fig 7). These HCT strains had large multiplications of the remaining part of the pathogenicity chromosome as well as co-transfer of contig 58 and part of contig 47 (Fig 8). These two contigs correspond to the accessory part of chromosome 3 and chromosome 6 of Fol4287, and they also correspond to the second transferred chromosome in Fol007 (4). Interestingly, for HCT_ΔGFP#8-1, HCT_ΔGFP#8-2 and HCT_ΔGFP#8-3, derived from ΔGFP#8 and HCT_ΔGFP#26-1, HCT_ΔGFP#26-2 and HCT_ΔGFP#26-3, derived from ΔGFP#26, all strains in which only a partial pathogenicity chromosome was present, higher virulence was also observed compared to HCT strains containing the complete pathogenicity chromosome (Fig 7 and Fig 8). In these cases, no extra sequences were co-transferred, and in ΔGFP#8-derived strains no multiplication of pathogenicity chromosome sequences was observed.

**Fig 7:**
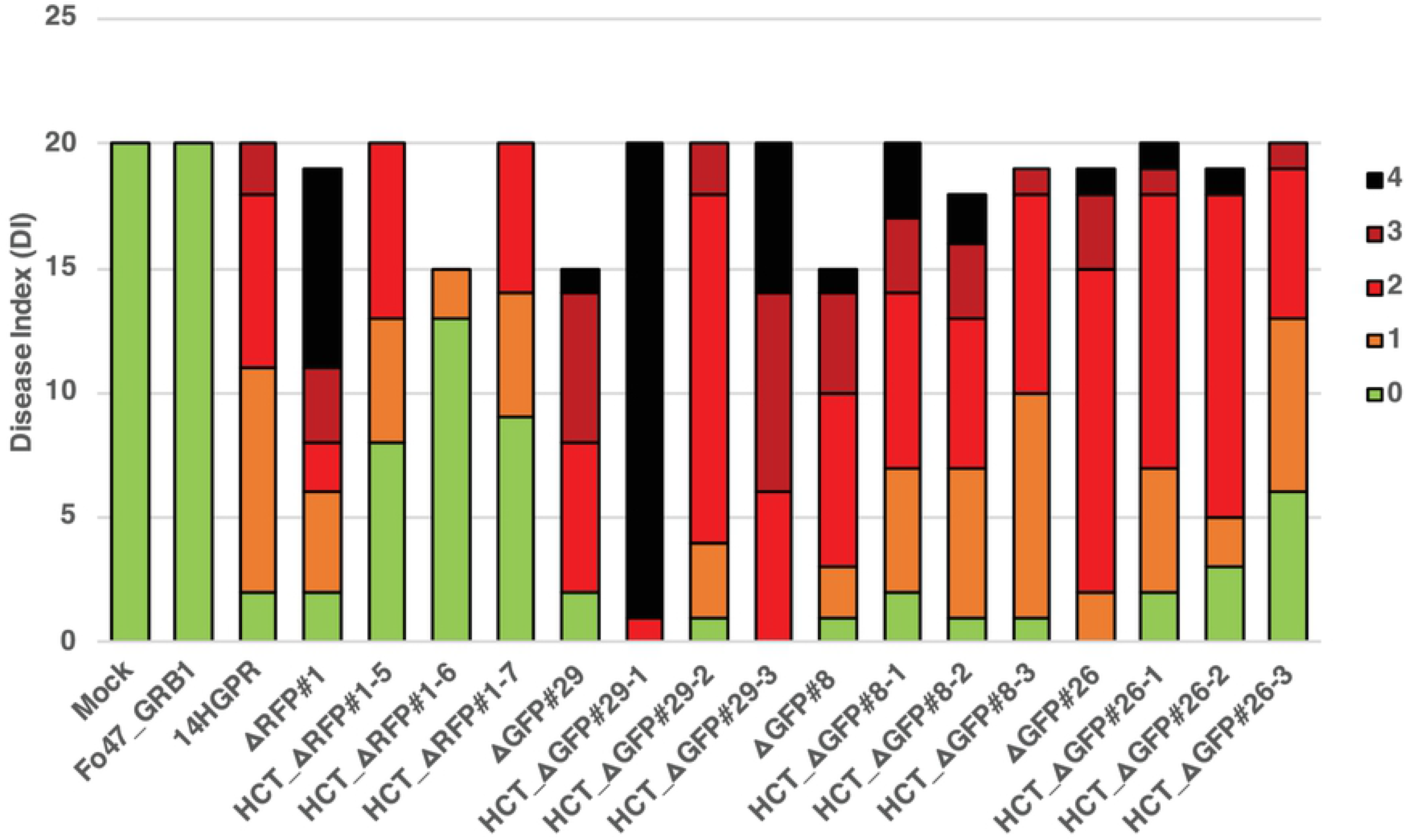
A partial pathogenicity chromosome can turn an endophyte into a pathogen. Bioassays were performed to assess the virulence of HCT strains. Ten days old tomato seedlings were inoculated with 1*10^7^ spores/mL at 25°C, and disease index (DI) of infected tomato plants was scored three weeks after inoculation. All four donor strains, ΔRFP#1, ΔGFP#29, ΔGFP#8 and ΔGFP#26 caused similar disease index compared to 14HGPR. Fo47_GRB1 did not cause any disease symptoms on tomato plants, while all HCT strains in the background of Fo47_GRB1 were able to cause disease on tomato plants, with some variation in Disease Index (DI). Disease index was scored on a scale of 0–4 (0, no symptoms; 1, one brown vessel below the cotyledons; 2, one or two brown vascular bundles at cotyledons; 3, three brown vascular bundles and growth distortion; 4, all vascular bundles are brown, plant either dead or very small and wilted). Kruskal-Wallis test was performed on disease index.

**Fig 8:**
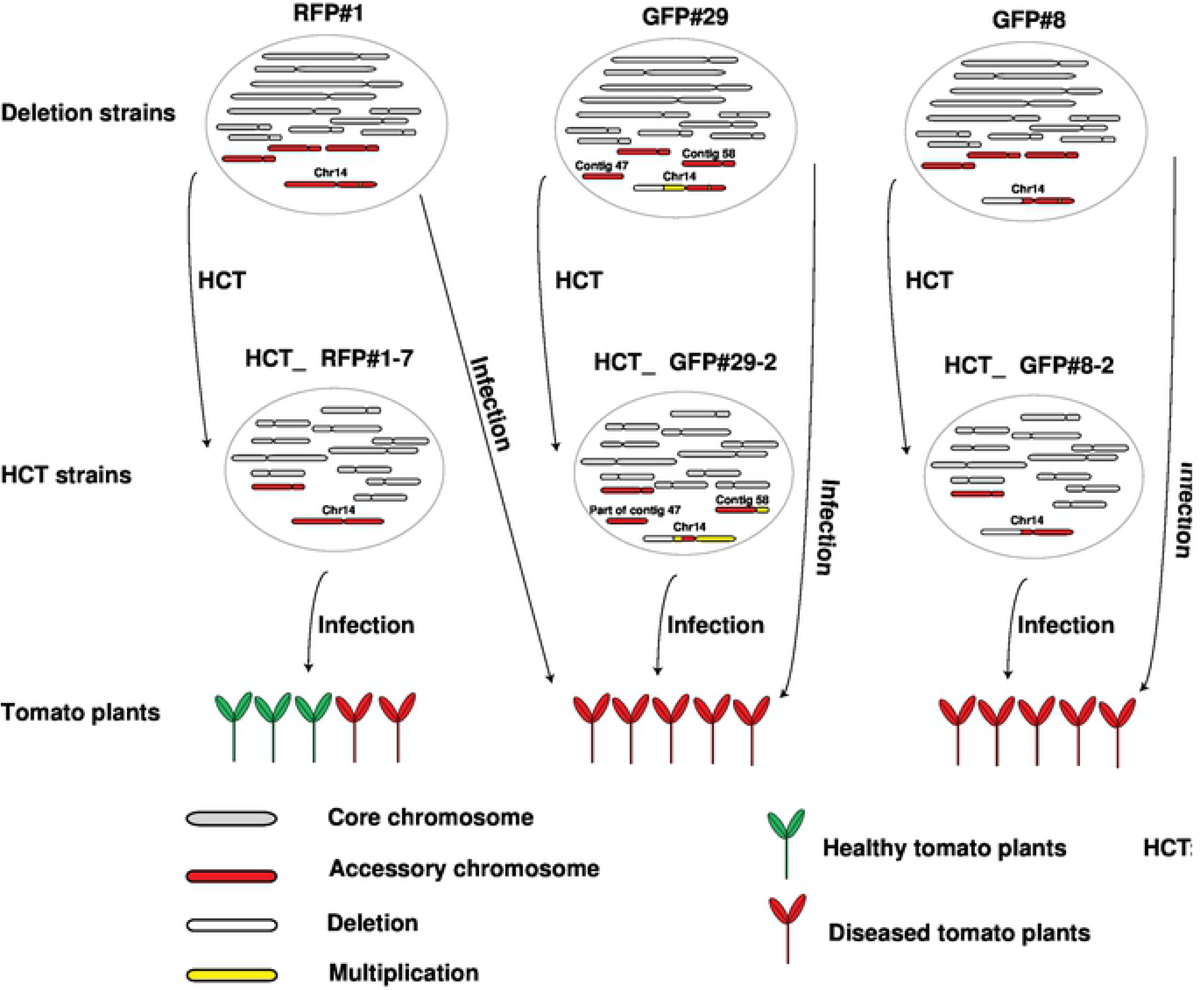
A partial pathogenicity chromosome in *Fusarium oxysporum* is sufficient to cause disease and can be horizontally transferred. Graphical summary of key observations. Strain ΔRFP#1 with a small deletion in the *RFP* region of the pathogenicity chromosome was still able to cause disease on tomato plants. In addition, it was able to transfer its pathogenicity chromosome to Fo47, turning the recipient strain into a tomato-infecting strain but with only mild virulence. Deletion strain ΔGFP#29 had lost a large part of the long (q) arm of the pathogenicity chromosome (white bar) and underwent a duplication of the remaining part of the q arm (yellow bar). This deletion strain still caused disease on tomato plants. Surprisingly, not only the partial pathogenicity chromosome but also (the rest of) contig 58 and part of contig 47 were transferred from ΔGFP#29 to Fo47. These transferred sequences most likely form one big chromosome, according to single chromosome sequencing (Fig S6) and CHEF gel analysis (Fig S5). The resulting HCT strain HCT_ΔGFP#29-2 caused severe disease on tomato plants. Deletion strain ΔGFP#8 had lost a large part of the q arm of the pathogenicity chromosome but it still caused disease on tomato plants. The partial pathogenicity chromosome in ΔGFP#8 could be transferred into Fo47, turning the recipient strain into a tomato pathogen (HCT_ΔGFP#8-2) with the same virulence as its donor, ΔGFP#8. Deletion strain ΔGFP#26 had completely lost the q arm of the pathogenicity chromosome and underwent an almost complete duplication of the short arm (p arm). This version of the pathogenicity chromosome could also be transferred into Fo47. Both the donor strain ΔGFP#26 and the recipient strain HCT_ΔGFP#26-1 caused disease on tomato plants to similar levels.

## Discussion

The pathogenicity chromosome in Fol is required for infecting tomato plants (4,16), and can be horizontally transferred to a non-pathogenic strain, turning the latter into a tomato pathogen (4,5). Here, we narrow down the regions and genes on the Fol pathogenicity chromosome that are required for virulence. Furthermore, we demonstrate that a partial pathogenicity chromosome can still be horizontally transferred to a non-pathogenic strain, and this is sufficient to turn that strain into a pathogen. Surprisingly, transfer of a partial pathogenicity chromosome leads to higher virulence than transfer of a complete pathogenicity chromosome. Possibly, sequences in the missing (q) arm suppress virulence in the genetic background of the non-pathogenic recipient strain, Fo47.

### How many effector genes does Fol need to infect its host?

Effector genes have been predicted and studied in many different plant pathogenic fungi (27–32), including *Cladosporium fulvum* (33–38), *Fusarium oxysporum* (19,19–21,39,40,40,41), *Leptosphaeria maculans* (42–48), *Magnaporthe oryzae* (49–54), *Melampsora lini* (31,55,56), and *Blumeria graminis* (57–59). In many cases, deletion of a single effector gene has little effect on virulence, suggesting functional redundancy of effectors (28). Nevertheless, it is likely that a limited number of effectors are required for virulence (5). In Fol, a single chromosome contains all effector genes (*SIX* genes) required and sufficient for infecting tomato (4,5,18). This provides the opportunity to study the minimal regions or genes on this chromosome that are sufficient for infection. Previously, several *SIX* genes have been shown to contribute to virulence, including *SIX1* (19), *SIX3* (20) and *SIX5* (21). Moreover, Vlaardingerbroek and coworkers (5) have shown that most of the long (q) arm of the pathogenicity chromosome, including *SIX9*, *SIX6* and *SIX11*, is not required for virulence. Here, we confirm that the complete q arm of the pathogenicity chromosome is dispensable for pathogenicity. We further demonstrate that a large part of the p arm is also not required for virulence. This part contains *SIX10/12/7* gene cluster, indicating that the in xylem secreted proteins Six7, Six10 and Six12 are not required for virulence. Deletion strains with a larger deletion of the p arm are not able to infect tomato plants, suggesting that the remaining part of the p arm of the pathogenicity chromosome is required for virulence. This region contains *SIX14*, *SIX1*, *SIX2*, *SIX3, SIX5*, and one copy of *SIX13*. We conclude that, although all *SIX* genes are highly expressed during infection (25) and the corresponding Six proteins are abundant in the xylem sap of infected tomato plants (18), only a subset of these proteins are required for virulence.

### Except effector genes, what else on the pathogenicity chromosome does Fol need to infect tomato?

It has been shown that a homolog of transcription factor gene *FTF1* (FOXG_17084), which is located close to *SIX2*, can induce expression of most *SIX* genes when overexpressed (25). Two additional homologs of *FTF1* are present on the pathogenicity chromosome – one is close to *SIX11* and another one is close to the *SIX10/12/7* gene cluster. Deletion strains without either of the latter two *FTF1* homologs are still fully virulent on tomato plants, indicating that they are dispensable for virulence. All deletion strains that are still virulent, however, contain the *FTF1* homolog FOXG_17084, so this homolog may be important for virulence (25).

In addition, a predicted secondary metabolite gene cluster, including seven genes, is located close to the *SIX10/12/7* gene cluster (18). This cluster can also be lost without affecting virulence. Besides *SIX14*, *SIX1*, *SIX2*, *SIX3, SIX5*, *SIX13* and the *FTF1* homolog FOXG_17084, approximately 70 additional predicted protein-coding genes reside within the part of the pathogenicity chromosome that all virulent deletion strains have in common, and some of these might contribute to virulence.

### How stable is the Fol genome?

The genomes of many plant pathogenic fungi have a high level of structural variation (8,60–64), including conserved core genome and lineage-specific regions or chromosomes that are characterized by a relatively high number of repetitive elements. The core genome is generally rather stable, but the lineage-specific regions are much more dynamic (5,11,16,65). In Fol strain 4287, lineage-specific regions include four accessory chromosomes (3, 6, 14 and 15), part of core chromosome 1 and part of core chromosome 2 (4). The differences in stability (loss and duplication) between the core genome and the accessory genome has been investigated previously (5). For example, the loss frequency of lineage-specific chromosomes was estimated to be approximately 1 in 35,000 in spores in a liquid culture. Surprisingly, core chromosome 12 in Fol4287 can also be lost but at a very low rate of 1 in 190,000 spores (5,66). In *Zymoseptoria tritici*, spontaneous accessory chromosome loss rate was much higher with chromosome loss in 2 to >50% of cells during four weeks of incubation (65). In the present study, we observed small deletions and duplications in core chromosomes, tending to occur at the end of chromosomes, as has also been observed in *Zymoseptoria tritici* (65). We also observed that (part of) the pathogenicity chromosome can be lost frequently (around 6 in 100,000 spores) (Fig 1B). It appeared that the p-arm of the pathogenicity chromosome is more stable than the q-arm (Fig 1B). However, this could be explained by multiplication of the region where RFP was inserted in the p-arm (Fig 2). Presumably, this region has undergone multiplication following homologous recombination in the original strain. To conclude, in Fol, we can confirm that the core genome is rather stable except for the telomeric regions, while the accessory chromosomes are relatively dynamic.

In the present study, a large variety of deletion and multiplication patterns have been observed for the pathogenicity chromosome, and almost all the deletions or multiplications have occurred in or close to repetitive elements. It is well known that repetitive elements can lead to intra- or inter-chromosome homologous recombination, resulting in deletions or translocations (67). It has also been shown that in some fungi the facultative heterochromatin mark H3K27me3 is present in both the subtelomeric regions of the core chromosomes and accessory chromosomes (12,66). This difference in histone modification compared to the core chromosomes may play a role in the difference in chromosome stability (66,68).

A highly dynamic accessory genome may accelerate the evolution of the pathogen in the arms race with its host (69). Many effector genes are located in the accessory part of the genome in many plant pathogenic fungi (4,8,70). However, effectors can be recognized by R proteins in plants, resulting in an immune response. Mutation or loss of effector genes can help to avoid recognition and regain virulence. The accessory part of the genome may provide a niche for rapid diversification of effector genes without influencing basic cellular functions.

## Materials and Methods

### Cloning

To replace *FOXG_14135*, *FOXG_16428* or *SIX10/12/7* with *RFP*, three constructs pRW1p_Pfem1_RFP_FOXG_14135, pRW1p_Pfem1_RFP_FOXG_16428, and pRW1p_Pfem1_RFP_SIX10/12/7 were made. Each of them contains a right border (facilitating *Agrobacterium tumefaciens* mediated transformation), the flanking sequences of each gene, the *FEM1* promoter, the *RFP* open reading frame (ORF), the *SIX1* terminator, the trpC terminator, the phleomycin ORF resistance cassette, the gpd promotor, another flank of each gene, and the left border. Firstly, pRW2h_Pfem_RFP_Tsix1 was constructed by amplifying the *RFP* ORF from pPK2-HPH-RFP (41) using primers FP6992 (AAAtctagaATGGCCTCCTCCGAGGACG) and FP6993 (TTTagatctTTAGGCGCCGGTGGAGTGG) followed by *XbaI*-*BglII* digestion and inserting it into the *XbaI*-*BglII* site of pRW2h_Pfem_MCS_Tsix1 (25). Then the hygromycine resistance cassette of pRW2h_Pfem_RFP_Tsix1 was replaced by the phleomycin resistance cassette of pRW1p_Pfem_MCS_Tsix1, which was modified from pRW1p (25,40). This resulted in pRW1p_Pfem_RFP_Tsix1. For the *FOXG_14135* deletion construct, around 1kb flanking regions of *FOXG_14135* were amplified using primers listed in Table S11. The two fragments were introduced into pRW1p_Pfem_RFP_Tsix1 using the HiFi cloning kit [New England Biolabs (UK) Ltd.]. The same method was used to make *FOXG_16428* and *SIX10/12/7* deletion constructs (Table S11). All constructs were checked by sequencing.

### Gene replacement in Fol

14HG6B was transformed via *Agrobacterium* mediated transformation (Table S1), as described previously (71). Transformants were monospored by pipetting 10 μl of sterile water on the emerging colony, and spreading this on a fresh Potato Dextrose Agar (PDA) plate supplemented with cefotaxime and Phleomycin. After two days of growth at 25°C, single colonies were picked and transferred to fresh plates. From these plates, glycerol stocks were made and these are the transformants we worked with.

### Fluorescence Assisted Cell Sorting (FACS)

Fluorescence Assisted Cell Sorting was used to select independent chromosome 14 deletion strains (24). Firstly, 14HGPR was mono-spored and single colonies were transferred to flasks with NO_3_ medium (0.17% yeast nitrogen base, 3% sucrose, 100mM KNO_3_) either directly or grown on PDA plates for some time before transferring to the NO_3_ medium. After growing for 5-7 days, spore suspensions were obtained by filtering cultures through a double layer of mira-cloth. To select spores without green or red fluorescence, 25 red (not green) and 25 green (not red) fluorescent spores were deflected on each plate and allowed to form colonies for 2-3 days at 25°C. The colonies were observed using the AMG Evos FL digital inverted microscope to confirm loss of red fluorescence or green fluorescence. Confirmed colonies were transferred to new plates and allowed to grow for at least two weeks before DNA extraction. To determine which parts of the pathogenicity chromosome (chromosome 14) were lost, PCR primers (Table S12) were used to scan the chromosome.

### Bioassays

To test virulence of Fol transformants, deletion strains or horizontal chromosome transfer strains on tomato (line C32), the root dip method was used (19). Briefly, spores were collected from 5-day-old cultures NO_3_ medium (0.17% yeast nitrogen base, 3% sucrose, 100mM KNO3) by filtering through miracloth (Merck; pore size of 22–25μm). Spores were centrifuged, resuspended in sterile MilliQ water, counted, brought to a final concentration of 1*10^7^ spores/mL and used for root inoculation of 10-day-old tomato seedlings. The seedlings were then potted individually and kept at 25 °C. Three weeks after inoculation, plant weight above the cotyledons was measured, and the extent of browning of vessels in the remaining part of the stem was scored. Disease index was scored on a scale of 0–4 (0, no symptoms; 1, one brown vessel below the cotyledons; 2, one or two brown vascular bundles at cotyledons; 3, three brown vascular bundles and growth distortion; 4, all vascular bundles are brown, plant either dead or very small and wilted).

### Horizontal chromosome transfer

To test whether partial pathogenicity chromosomes can be transferred or not, horizontal chromosome transfer experiments were performed (26). In total, 24 deletion strains (Table S8) were selected to co-incubate with Fo47pGRB1 (17) or Fo47-H1 (4). Strains were grown in minimal liquid medium (3% sucrose, 0.17% yeast nitrogen base and 100mM KNO_3_) for 3-5 days, after which 10^5^ or 2×10^5^ microconidia from the donor and recipient strains were mixed in different ratios and co-incubated on PDA or Czapek Dox Agar (CDA) plates for eight days. Spores were collected from these plates using 2-5 ml sterile MilliQ, filtered through sterile miracloth and pipetted on a double selective PDA plate containing 0.1 M Tris pH 8 supplemented with 100 μg/ml hygromycin (Duchefa) and 100 μg/ml zeocin (InvivoGen). Double drug resistant colonies were selected after three days and monospored on a new plate supplemented with both drugs. After two to three days of growth, colonies were selected and transferred to new plates supplemented with zeocin and hygromycin. Fluorescence of double drug-resistant colonies was checked with an AMG Evos FL digital inverted microscope. Strains with both red and green fluorescence were allowed to grow for 2 weeks before DNA isolation. Both selection markers and other genes (Table S10) were used to confirm horizontal chromosome transfer by PCR.

### Contour-clamped homogeneous electric field (CHEF) electrophoresis

To confirm horizontal chromosome transfer, Contour-clamped homogeneous electric field (CHEF) electrophoresis was performed. Preparation of protoplasts and pulsed-field gel electrophoresis were performed as described previously (4). *Fusarium* strains were cultured in 100 ml NO_3_ medium (0.17% yeast nitrogen base, 100 mM KNO_3_ and 3% sucrose) for five days at 25 °C. Then, conidia were collected by filtering through a double-layer of miracloth and the concentration of spores were measured. Five × 10^8^ conidia were transferred to 40 ml PDB (BD biosciences). After approximately 16 hours of growth at 25 °C, germinated spores were re-suspended in 10 ml MgSO_4_ solution (1.2 M MgSO_4_, 50 mM sodium citrate, pH 5.8) supplemented with 50 mg/ml Glucanex (Sigma) + 5 mg/ml driselase (Sigma, D9515) and incubated for approximately 17 hours at 30°C in a shaking water bath (65 rpm). The protoplasts were filtered through a double layer of miracloth, collected by centrifugation and casted in InCert agarose (Lonza) at a concentration of 2 × 10^8^ protoplasts per ml. Plugs were treated with 2 mg/ml pronase E at 50°C. Chromosomes were separated by pulsed-field electrophoresis for 260 hours in 1% Seakem Gold agarose (Lonza) at 1.5 V/cm in a CHEF-DRII system (Biorad) in 0.5 × TBE at 4 °C, with switch times between 1200 and 4800 s. The gels were stained with 1μg/mL ethidium bromide in 0.5 × TBE.

### Single chromosomes recovery from a CHEF gel

Chromosome DNA recovery from CHEF gels were performed according to the method described previously (72). Chromosome bands of interest were excised from the gel and were placed in 2 ml microcentrifuge tubes, then heated at 100°C while shaking at 350 rpm for at least 10 minutes to melt the agarose. After the melting step, six units of thermostable β-agarase (Nippon gene, Tokyo, Japan) were added to the gel solution, and held at 57°C, 350 rpm for 15 min. After enzyme treatment, tubes were kept on ice for 15 min to confirm the agarose was completely digested. If remaining agarose was observed in the reaction mixture, melting (100°C for 10 min) and subsequent steps were repeated. The concentration of DNA in the reaction mixture was checked by a Qubit 3.0 fluorometer (Invitrogen, Carlsbad, CA, USA) and the Qubit dsDNA HS Assay kit (Invitrogen).

### Illumina single chromosome and whole genome sequencing

Genomic DNA isolation was performed on freeze-dried mycelium ground in liquid nitrogen as starting material, using multiple rounds of phenol-chloroform extraction and precipitation, as well as the Purelink plant total DNA purification kit (Invitrogen).

Illumina sequencing (150 bp paired-end, insert size ~500 bp) was performed on a HiSeq 2500 machine at the Hartwig Medical Foundation (Amsterdam, the Netherlands) at ~100X coverage, resulting in 5.0–5.6 Mb of sequence data per sample.

Raw reads were trimmed to remove low-quality bases and adapter sequences using fastq-mcf v1.04.807 (-q 20). PCR duplicates were removed using PicardTools MarkDuplicates v2.7.1 with standard settings.

To assess partial deletions of the pathogenicity chromosome, reads of deletion strains were mapped directly to the SMRT assembly of Fol4287.

Reads from single chromosomes were also mapped directly to the SMRT assembly of Fol4287. To confirm horizontal chromosome transfer, trimmed reads were directly mapped to SMRT assembly of Fol4287, and only reads that mapped once with 100% coverage and 100% identity were selected (with SAMtools view ‒q 42) when calculating read densities.

For visualization of the reads counts in 10 kb non-overlapping sliding windows, SAMtools bedcov was used. SAMtools version 1.8 was used in all above-mentioned cases.

## Acknowledgements

We are grateful to Petra Houterman for the help with CHEF gel experiments; Harold Lemereis and Ludek Tikovsky for plant care; J.L. was financially supported by the China Scholarship Council program (File number: 201504910768). L.F. was financially supported by the NWO Talent Scheme Veni (Grant number: 016.veni.181.090). No conflict of interest is declared.

## Supporting information

**Fig S1: Deletion of *FOXG_14135* does not result in reduced virulence.**

Fresh weight (A) and disease index (DI) (B) of infected tomato plants were scored three weeks after inoculation. When ten days old tomato seedlings were inoculated with 1*10^7^ spores/mL at 25°C, the *FOXG_14135* deletion strain 14HGPR showed similar disease index and fresh weight as the original strain 14HG6B. As control, disease symptoms of an ectopic transformant (T-DNA was randomly inserted in the genome) were assessed, and no significant difference in virulence was observed compared to 14HG6B. Water (Mock)-treated plants were completely healthy. Disease index was scored on a scale of 0–4 (0, no symptoms; 1, one brown vessel below the cotyledons; 2, one or two brown vascular bundles at cotyledons; 3, three brown vascular bundles and growth distortion; 4, all vascular bundles are brown, plant either dead or very small and wilted). One-way ANOVA was performed on fresh weight. Kruskal-Wallis test was performed on disease index.

**Fig S2: Illumina whole genome read mapping of *GFP* deletion strains reveals a few minor changes in the core genome.**

(A) Whole genome reads of nine *GFP* deletion strains were mapped to the SMRT assembly of Fol4287. As reference, Illumina reads of Fol4287 itself were also mapped. For comparison of differences within and between deletion strains, genome coverage was normalized. No obvious changes were observed in the core genome of the three previously generated strains 14-4, 14-7, and 14-2. For all the deletion strains generated in this study, the same small deletion was observed at the end of contig 0. In addition, ΔGFP#20, ΔGFP#22, ΔGFP#8, and ΔGFP#29 all showed the same deletion in contig 47. For ΔGFP#29, deletions at the end of contig 3 and contig 7 and one duplication at the end of contig 61 were also observed. GC content (B) and repeat distribution across the genome (C) are also displayed.

**Fig S3: Illumina whole genome read mapping of *RFP* deletion strains reveals a few minor changes in the core genome.**

(A) Whole genome reads of four *RFP* deletion strains were mapped to the SMRT assembly of Fol4287. As reference, Illumina reads of Fol4287 itself were also mapped. For comparison of differences within and between deletion strains, genome coverage was normalized. For all the deletion strains, the same small deletion was observed at the end of contig 0 and in the middle of contig 47. For ΔRFP#12, a deletion at the end of contig 2 was observed. GC content (B) and repeat distribution across the genome (C) are also displayed.

**Fig S4: Stringent selection of mapped Illumina reads of HCT strains and donor strains to the SMRT assembly of Fol4287 shows absence of core chromosome transfer and confirms transfer of accessory regions.**

(A) Illumina reads of HCT strains (HCT_ΔGFP#29-2, HCT_ΔGFP#8-2 and HCT_ΔGFP#26-1) and their respective donor strains (ΔGFP#29, ΔGFP#8 and ΔGFP#26-1) were mapped to the SMRT assembly of Fol4287, and only those reads that mapped completely and without any mismatches were selected. In the case of transfer of core chromosomes, a high density of perfectly mapped reads was expected, even in the sub-telomeric regions as shown for the reference donor strains. No core chromosome transfer was observed for any HCT strain. (Partial) pathogenicity chromosome transfer was confirmed for all HCT strains. In HCT_ΔGFP#29-2, co-transfer of contig 58 and part of contig 47 was observed. GC content (B) and repeat distribution across the genome (C) are also displayed.

**Fig S5: Single chromosomes cut from a CHEF gel for sequencing.**

Eight strains were selected for chromosome separation in a CHEF gel. In total, 11 bands were cut from the gel (1 to 11) and sent for sequencing. The numbers are indicated on the respective bands. The same number indicates corresponding bands from the same strain. The name used in the main text for each number is listed below the gel. Chromosomes of *S. cerevisiae* were used as marker.

**Fig S6: Single chromosome sequencing confirms partial pathogenicity chromosomes in both donor and HCT strains.**

(A) Illumina reads of 11 bands cut from a CHEF gel (see Fig. S5) were mapped to the SMRT assembly of Fol4287. As reference, Illumina reads of Fol4287 itself were also mapped. Except band ΔGFP#29_SC_XS, which was not successfully sequenced, the remaining ten bands indeed contained sequences of the pathogenicity chromosome (contig 14). For example, the same partial pathogenicity chromosome is present in donor strains (ΔGFP#8 and ΔGFP#26) and the respective recipient strains (HCT_ΔGFP#8-2 and HCT_ΔGFP#26-1). Band ΔGFP#29_SC_S contained sequences from contig 7 and part of the pathogenicity chromosome. Instead of band HCT_ΔGFP#29_SC_S, band HCT_ΔGFP#29_SC_L was confirmed to be the transferred chromosome, containing sequences of the pathogenicity chromosome as well as contig 58 and part of contig 47. GC content (B) and repeat distribution across the genome (C) are also displayed.

**Table S1: Summary of *Agrobacterium*-mediated *Fusarium* transformations.**

**Table S2: Fol pathogenicity chromosome deletion strains obtained.**

Symbols used in the table: + for positive PCR result, − for negative PCR result; grey regions without symbol for presumed presence, white regions without symbols for presumed absence.

**Table S3. Summary of three Fluorescence Assisted Cell Sorting (FACS) experiments.**

**Table S4. Details of the second Fluorescence Assisted Cell Sorting (FACS) experiment.**

**Table S5. Details of the third Fluorescence Assisted Cell Sorting (FACS) experiment.**

**Table S6. Details of the fourth Fluorescence Assisted Cell Sorting (FACS) experiment.**

**Table S7: Strains and single chromosomes sent for sequencing.**

**Table S8. Summary of Horizontal Chromosome Transfer (HCT) experiments.**

No: no successful transfer; Yes: successful transfer. Only strains for which transfer was attempted are shown in this table.

**Table S9. Different strain ratios and media used in five HCT experiments.**

PDA: potato dextrose agar; CDA: Czapek Dox Agar. CAT medium: 0.17% YNB, 25 mM KNO_3._

**Table S10. Horizontal chromosome transfer was confirmed by PCR.**

Symbols used in the table: + for positive PCR result, − for negative PCR result; black regions without symbol are presumed to be present, white regions without symbols are presumed to be absent.

**Table S11: Primers used for cloning.**

**Table S12: Markers on the pathogenicity chromosome.**

